# The role of the endoplasmic reticulum in *in vivo* cancer FDG kinetics

**DOI:** 10.1101/664417

**Authors:** Mara Scussolini, Vanessa Cossu, Cecilia Marini, Gianmario Sambuceti, Giacomo Caviglia, Michele Piana

## Abstract

A very recent result obtained by means of an *in vitro* experiment with cancer cultured cells has configured the endoplasmic reticulum as the preferential site for the accumulation of 2-deoxy-2-[^18^F]fluoro-D-glucose (FDG). Such a result is coherent with cell biochemistry and is made more significant by the fact that reticular accumulation rate of FDG is dependent upon extracellular glucose availability. The objective of the present paper was to confirm this result *in vivo*, using small animal models of CT26 cancer tissues. Specifically, assuming that the endoplasmic reticulum plays a specific functional role in the framework of a three-compartment model for FDG kinetics, we are able to explain positron emission tomography dynamic data in a more reliable way than by means of a standard Sokoloff two-compartment system. This result is made more solid from a computational viewpoint by means of some identifiability considerations based on a mathematical analysis of the compartmental equations.

## 1 INTRODUCTION

2-deoxy-2-[^18^F]fluoro-D-glucose (FDG) is widely used as a glucose analogue tracer to evaluate glucose metabolism in living organisms. As glucose, FDG is first transported into cells, where it is phosphorylated to FDG-6-phosphate (FDG6P) by hexokinase (HK) and then accumulates intracellularly. The measured amount of emitted radiation is considered an accurate marker of overall glucose uptake by cells and tissues (Cherry et al., 2012). In addition, FDG consumption by cancer cells is increased by the Warburg effect for glucose (Vander et al., 2009); consequently, FDG may be employed in cancer detection and staging, and to determine the effectiveness of medical treatments (Cairns et al., 2011).

A recent result concerning the characterization of FDG kinetics in cultured cancer cells (Scussolini et al., 2019) showed that the endoplasmic reticulum (ER) is the preferential site of FDG accumulation and that, even more importantly, FDG reticular accumulation rate is dependent upon extracellular glucose availability. This result is coherent with recent progress in cell biochemistry, which clarified that the glucose-6-phosphatase (G6Pase), responsible for the dephosphorylation, is compartmentalized within the endoplasmic reticulum (ER) (Ghosh et al., 2002; Marini et al., 2016).

The investigation in (Scussolini et al., 2019) relied on two methodological tools, one experimental and one computational. In fact, on the one hand, FDG kinetics in cells cultured over a Petri dish, was evaluated using the dedicated Ligand Tracer device (Björke and Andersson, 2006), which is able to count electron/positron events without contaminating the counting rate of the cultured cells. On the other hand, the data analysis was performed within the framework of a novel compartmental model, which extended the traditional Sokoloff two-compartment analysis (Sokoloff et al., 1977) by constraining G6Pase sequestration within the ER lumen.

However, this methodological approach left two significant issues open, again one experimental and one computational, that should be addressed to provide further significance to this biochemical finding. The first issue requires the *in vivo* confirmation of the crucial role that the FDG - ER connection plays in the *in vitro* metabolism of cancer cells. The second issue is concerned with the mathematical identifiability of the three-compartment model utilized to prove this connection, which must be discussed in order to make the model sound and not ambiguous from the data analysis perspective.

The main objective of the present paper is to show the central role of ER for the FDG kinetic in *in vivo* tissues. This result is obtained by processing FDG-PET data of murine models inoculated with specific murine cancer cell lines through a three-compartment model for FDG concentration. This model is the analog of the one utilized in (Scussolini et al., 2019) for describing the FDG kinetic in the case of *in vitro* cultured data and is designed according to the biochemically-driven assumption that most FDG is phosphorylated in ER. This assumption is confirmed by the experimental results obtained by using six datasets provided by a PET scanner for small animals in the case of six murine models of CT26 colon cancer. Further, the reliability of the results is corroborated by some identifiability considerations based on a formal and numerical analysis of the compartmental equations.

The structure of the paper is as follows. Section 2 recalls the three-compartment model utilized for the analysis. Section 3 provides a discussion the model identifiability. Section 4 describes the application of the model to the *in vivo* data and Section 5 discusses the results by comparing them with the ones obtained by using a standard two-compartment model. Our conclusions are offered in Section 6.

## 2 THE COMPARTMENTAL MODEL

The three-compartment model introduced in (Scussolini et al., 2019) and utilized in the present paper relies on the biochemical considerations schematically illustrated in Figure 1. Specifically, in this scheme FDG is transported through the cell membrane by GLUT proteins. Inside cytosol, FDG is phosphorylated by HK to FDG6P. Unlike G6P, FDG6P is a false substrate for downstream enzymes channeling G6P to glycolysis (G6P-isomerase) or pentose-phosphate pathway (G6P-dehydrogenase). However, FDG6P is a substrate for the enzyme G6Pase and thus it can be dephosphorylated. Recent advances in biochemistry show that G6Pase is anchored to the ER (Ghosh et al., 2002), so that its action of hydrolysis of FDG6P, resulting in the creation of a phosphate group and free molecules, occurs after transport of the phosphorylated forms into the ER by the transmembrane protein glucose-6-phosphate transporter (G6PT) (Marini et al., 2016). Subsequently, FDG may be released in cytosol. Further biochemical, pharmacological, clinical, and genetic data lead to a natural interpretation of ER as a distinct metabolic compartment (Csala et al., 2007). This role of ER in FDG kinetics is extended to tissues by means of the biochemically driven three-compartment model (BCM from now on) illustrated in Figure 1, upper right panel. In this figure, BCM is compared to the traditional two-compartment Sokoloff model (SCM from now on). The compartments in BCM identified by the indexes *i*, *f*, *p*, and *r* refer to free tracer in the input pool, free tracer in cytosol, phosphorylated tracer in cytosol, and phosphorylated tracer in ER, respectively. In principle, ER may interchange free FDG molecules with the cytosol, so that free tracer can be found in ER; here we assume that the free tracer in ER reaches equilibrium almost instantaneously and represents a small fraction of the tracer contained in ER, so that input of free tracer from cytosol and the content of free tracer in ER are discarded.

**Figure 1.**
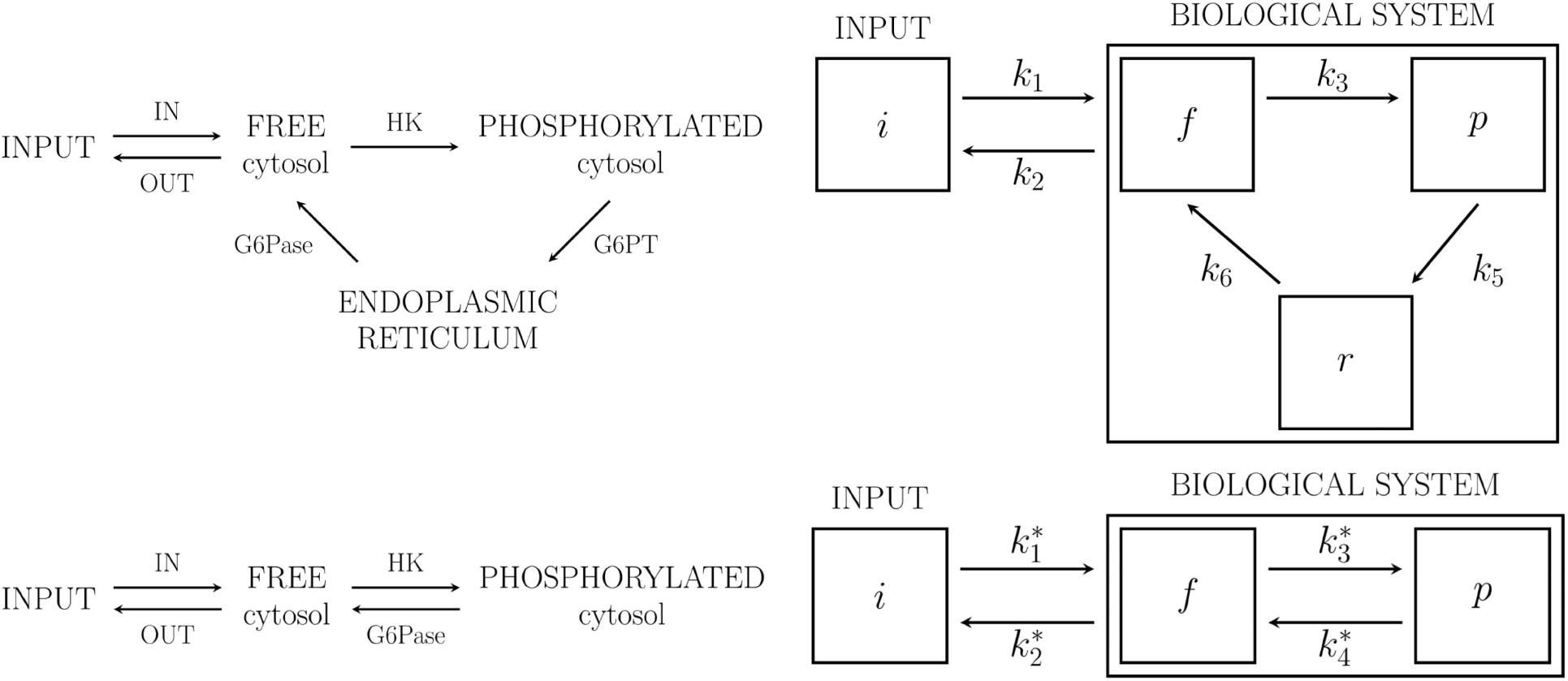
The compartmental models used for data analysis. Left upper panel: the FDG biochemical path assuming ER as accumulation site for FDG; right upper panel: corresponding biochemically-driven three-compartment model (BCM). Left bottom panel: the traditional Sokoloff path for FDG; the corresponding two-compartment Sokoloff model (SCM).

The rate constants *k*_*i*_ (1/min), with *i* ϵ {1, 2, 3, 5, 6}, describe the first order process of tracer transfer between compartments. According to Figure 1, upper right panel, *k*_1_ and *k*_2_ are the rate constants for transport of FDG from blood to tissue and back from tissue to blood (by GLUTs), respectively; *k*_3_ is the phosphorylation rate of FDG (by HK); *k*_5_ is the input rate of FDG6P into ER (by G6PT); *k*_6_ refers to the dephosphorylation rate of FDG6P to FDG (by G6Pase). Since dephosphorylation occurs only inside ER, a parameter *k*_4_, corresponding to an arrow from *p* to *f*, is not considered.

We assume that standard conditions for applicability of compartmental models are satisfied. In particular, the distribution of tracer in each compartment is spatially homogeneous, and tracer exchanged between compartments is instantaneously mixed (Cherry et al., 2012). We also assume that appropriate corrections for the physical decay of radioactivity have been applied.

The time-dependent functions *C*_*f*_, *C*_*p*_, and *C*_*r*_ represent the tracer concentrations of the model compartments, and are the state variables of the compartmental system. The tracer concentration of the input compartment *C*_*i*_ is considered as the given input function (IF). The Ordinary Differential Equations (ODEs) for the tracer flow are

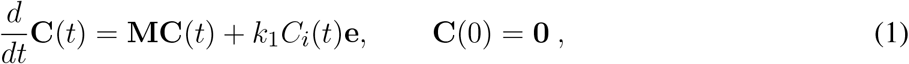

where

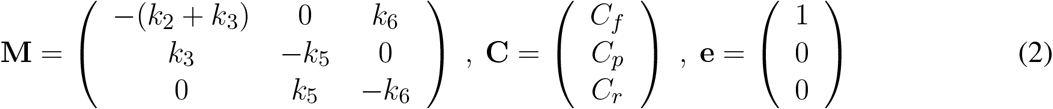

and with the time variable *t* ∈ ℝ_*>*0_. The initial condition **C**(0) = **0** means that no tracer amount is in the system at the beginning of the experiment. The analytic solution of the Cauchy problem (1) is

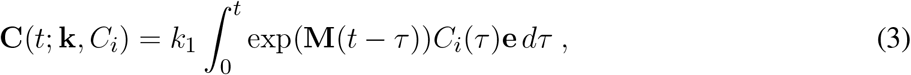

with 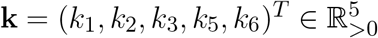.

*In vivo* experiments utilize sequences of PET images acquired at different time intervals. In each PET frame, two regions of interest (ROIs) are drawn, one highlighting the tumor and the other one identifying the left ventricle. The volume *V*_tot_ of the tumor ROI is partitioned as

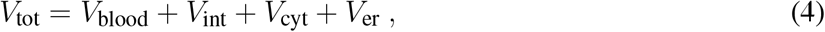

where *V*_blood_ and *V*_int_ denote the volume occupied by blood and interstitial fluid, respectively; *V*_cyt_ and *V*_er_ denote the total volumes of cytosol and ER in tissue cells. The total activity *V*_tot_𝒞_*T*_ in *V*_tot_ is related to the state variables and the IF by

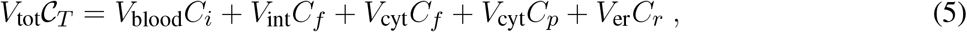

or, equivalently,

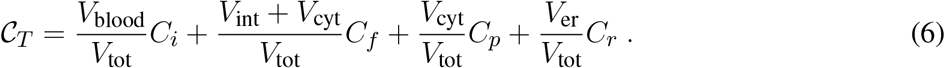

By defining the volume fractions of blood and interstitial fluid as

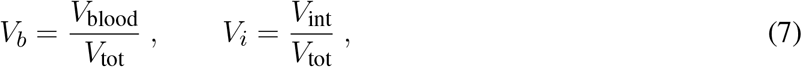

it is straightforward to obtain

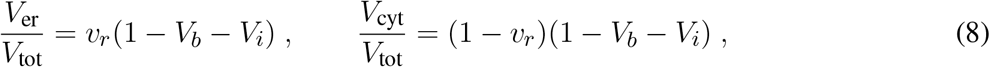

where

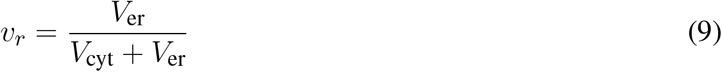

is independent of the number of cells. Replacement of (7) and (8) into (6) yields the required result:

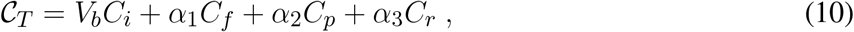

where the positive adimensional constants *α*_1_, *α*_2_, and *α*_3_ are defined as

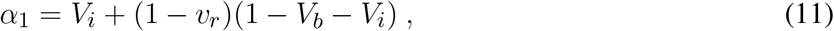

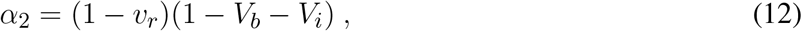

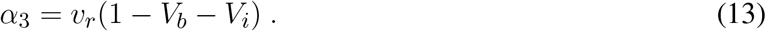

In the experimental applications of Section 4 we set *V*_*b*_ = 0.15 according to (Montet et al., 2007) and *V*_*i*_ = 0.3 according to (Kim et al., 2004); the volume fraction occupied by ER with respect to cytosol has been estimated as *v*_*r*_ = 0.14. In compact form, (10) becomes

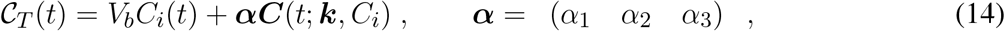

which is the inverse problem associated to BCM. In order to solve it, our approach followed a regularized Newton-type method (Bauer et al., 2009; Delbary and Garbarino, 2016), already validated and applied successfully in other compartmental problems, e.g. models for complex physiologies (Garbarino et al., 2014, 2015), parametric imaging (Scussolini et al., 2017), and reference tissue approaches (Scussolini et al., 2018). The algorithm is denoted as reg-GN in the following.

## 3 IDENTIFIABILITY ISSUES

Identifiability of linear compartmental models has been widely analyzed and there are a lot of results already available in the literature, some of which are collected in Cobelli et al. (2002). In particular, BCM as described by Figure 1, upper right panel, corresponds to model number 6 in Figure 5.7.2 at page 141 of that book, which provides a result of global identifiability. However, that result does not account for the fact that in experiments relying on PET images, the only available measurements correspond to the sum of the tracer concentrations in the left ventricle and in the tumor, as illustrated in equation (10). Nevertheless, the standard techniques illustrated in (Scussolini et al., 2017; Delbary et al., 2016) and relying on the use of the Laplace transform straightforwardly leads to the following result: assume that the polynomials

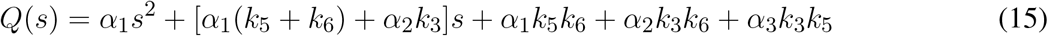

and

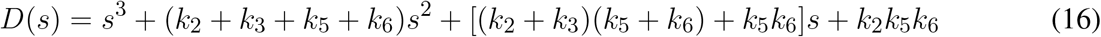

are coprime; if 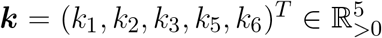 is generic, then ***k*** is uniquely determined by *C*_*i*_ and *𝒞*_*T*_, and the BC model of equations (1)–(10) is identifiable.

**Figure 2.**
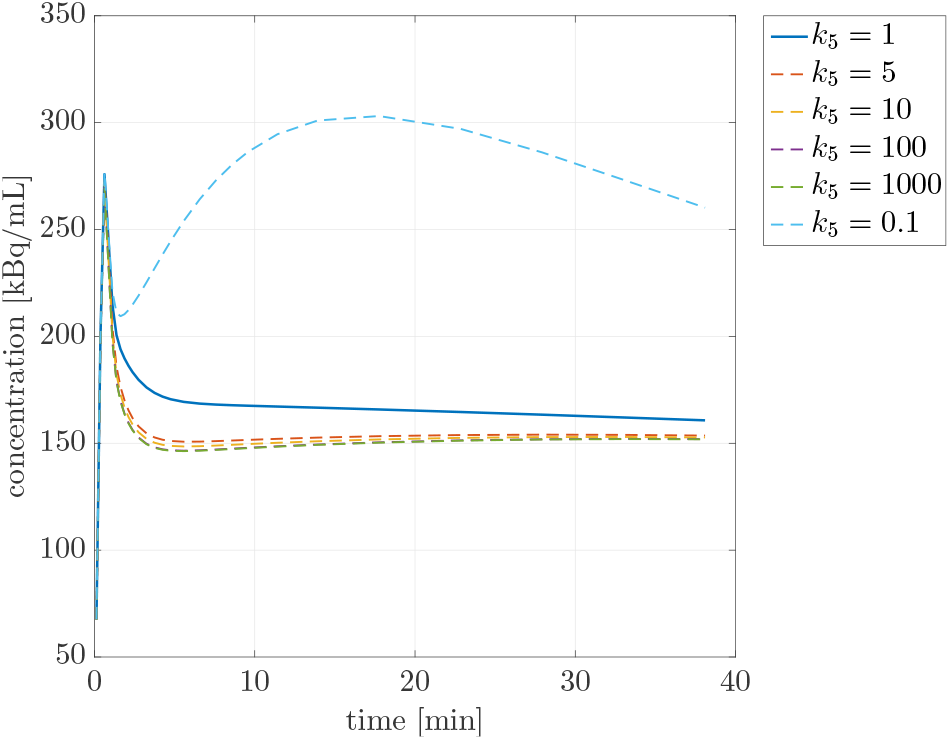
Sensitivity of BCM with respect to *k*_5_: plots of *C*_*T*_(*t*) provided by the model for 6 values of the kinetic parameter.

This formal result does not discard the possibility of numerical non-uniqueness and therefore of the unreliability of this compartmental analysis. In fact, a straightforward sensitivity analysis (Goulet, 2016) shows that the local sensitivity of *C*_*T*_ versus *k*_5_ is more than one order of magnitude lower than the other kinetic parameters. This ambiguity is clearly shown in Figure 2, where the forward problem (3) is computed for different values of *k*_5_ and the total concentration *C*_*T*_ is illustrated as a function of time: the plots in the figure points out that, for high values of *k*_5_, *C*_*T*_ remains almost constant even when *k*_5_ increases of three orders of magnitude. In order to account for this aspect in the data analysis process, the application of reg-GN can be realized either by initializing *k*_5_ randomly in (0, 1) or according to a simple heuristic procedure: first, perform an analysis of the same dataset using the standard SCM, which provides a first estimate 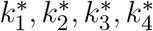 of its four kinetic parameters; then, initialize *k*_5_ with a value smaller than 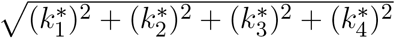. This initialization condition implements an energetic constraint on *k*_5_, which prevents it to reach ranges of values where the sensitivity of *C*_*T*_ becomes too small.

## 4 APPLICATIONS TO *IN VIVO* FDG-PET DATA

We analyzed a group of six mice, denoted as m*i*, with *i* = 1, …, 6, whose basic characteristics are reported in Table 1: cell-line type, sex, weight, glycemia, the peak value of the arterial IF *Ĉ*_*i*_, reached in the first few minutes of the PET acquisition, and the tracer concentration in the ROI tissue at the end time *𝒞*_*T*_(*t*_*f*_). The experimental ROI concentration *C*_*T*_ of the tumor for one of the mice (specifically, mouse m1, Figure 3, left panel) is shown in Figure 3, middle panel; the related canonical arterial input function *C*_*i*_ is shown in Figure 3, right panel.

**Table 1.** Experimental values for the FDG-PET measurements: type of cell line, sex, weight (g), and glycemia (mg/dL) for each mouse, maximum value of the IF *Ĉ*_*i*_ (kBq/mL), and final-time (*t*_*f*_) total concentration of the cancer tissue *𝒞*_*T*_(*t*_*f*_) (kBq/mL).

| | Cell type | Sex | Weight [g] | Glycemia [mg/dL] | $\hat{C}_i$ [kBq/mL] | $C_T(t_f)$ [kBq/mL] |
| --- | --- | --- | --- | --- | --- | --- |
| m1 | CT26 | *F* | 18.7 | 112 | $1.52 \cdot 10^3$ | $3.18 \cdot 10^2$ |
| m2 | CT26 | *F* | 17.2 | 84 | $1.91 \cdot 10^3$ | $2.73 \cdot 10^2$ |
| m3 | CT26 | *F* | 19.9 | 162 | $9.58 \cdot 10^2$ | $1.62 \cdot 10^2$ |
| m4 | CT26 | *F* | 16.8 | 81 | $1.54 \cdot 10^3$ | $3.53 \cdot 10^2$ |
| m5 | CT26 | *F* | 17.3 | 69 | $6.69 \cdot 10^2$ | $3.44 \cdot 10^2$ |
| m6 | CT26 | *F* | 15.9 | 53 | $1.34 \cdot 10^3$ | $3.01 \cdot 10^2$ |

**Figure 3.**
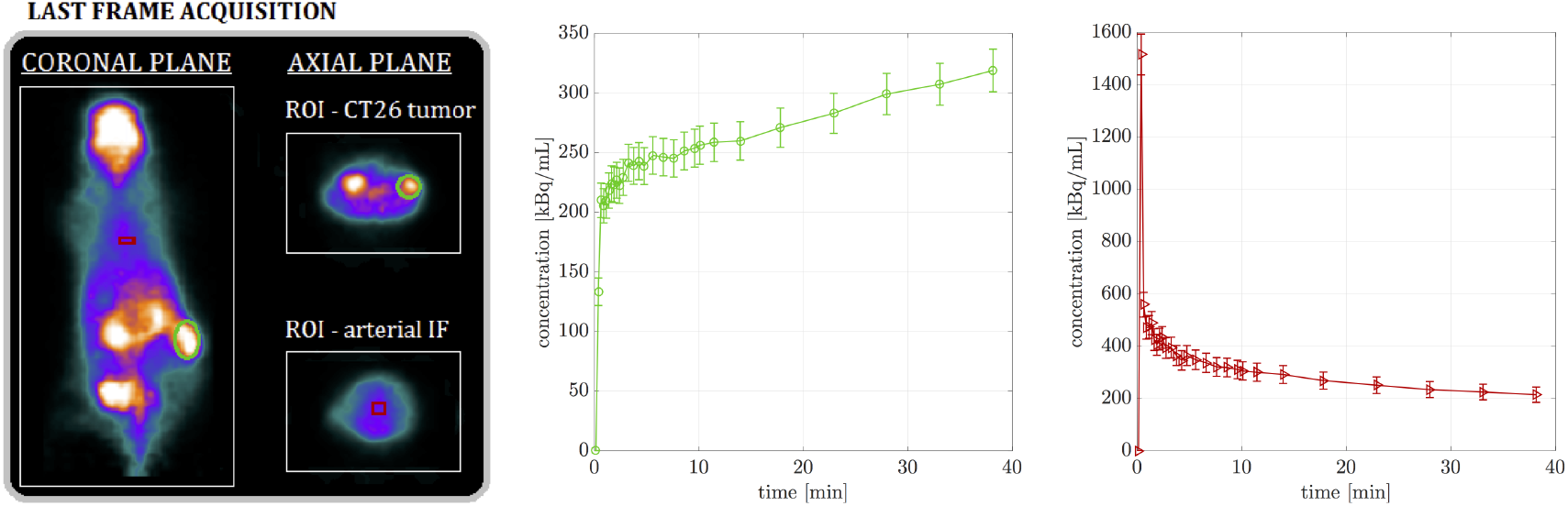
(a) Last frame of the FDG-PET acquisition of the murine model m1 with ROIs around the CT26 tumor (green color) and the aortic arc (red color). (b) The time-dependent ROI concentration curve of the CT26 tumor *𝒞*_*T*_ and its standard deviation, related to experiment m1. (c) The time-dependent concentration curve of the arterial input function *C*_*i*_ and its standard deviation, related to experiment m1.

FDG-PET data of CT26 cancer tissues have been processed by the application of both BCM and SCM. The numerical reduction of both models has been performed by means of reg-GN. Estimates of the parameters obtained for each experiment of the group of mice are reported in Table 2 for BCM, and in Table 3 for SCM. Means and standard deviations have been computed by repeating fifty runs of the reg-GN code, with fifty different random initialization values of the rate constants in the interval (0,1). The regularization parameter was determined at each iteration via Generalized Cross Validation method (Golub et al., 1979), with a confidence interval ranging between 10^4^ and 10^6^. The threshold for the stopping criterion of the iterative algorithm has been chosen of the order of 10^*−*1^, for both models.

**Table 2.** Reconstructed kinetic parameters (1/min) by the use of BCM for the CT26 tumor tissue of the FDG–PET experimental group of six mice, as mean and standard deviation over 50 runs of the reg-GN algorithm. The last two lines report mean and standard deviation of each kinetic parameter computed over the mean estimates of the six murine experiments.

| | $k_1$ | $k_2$ | $k_3$ | $k_5$ | $k_6$ |
| --- | --- | --- | --- | --- | --- |
| m1 | $0.32 \pm 0.03$ | $0.37 \pm 0.15$ | $0.45 \pm 0.19$ | $0.51 \pm 0.28$ | $0.03 \pm 0.02$ |
| m2 | $0.47 \pm 0.04$ | $0.67 \pm 0.14$ | $0.54 \pm 0.16$ | $0.59 \pm 0.26$ | $0.03 \pm 0.04$ |
| m3 | $0.17 \pm 0.03$ | $0.34 \pm 0.17$ | $0.58 \pm 0.21$ | $0.56 \pm 0.25$ | $0.03 \pm 0.02$ |
| m4 | $0.25 \pm 0.03$ | $0.22 \pm 0.12$ | $0.64 \pm 0.21$ | $0.58 \pm 0.23$ | $0.08 \pm 0.02$ |
| m5 | $0.30 \pm 0.04$ | $0.33 \pm 0.19$ | $0.85 \pm 0.31$ | $0.61 \pm 0.27$ | $0.09 \pm 0.05$ |
| m6 | $0.31 \pm 0.03$ | $0.36 \pm 0.12$ | $0.61 \pm 0.21$ | $0.53 \pm 0.27$ | $0.09 \pm 0.03$ |
| mean | 0.30 | 0.38 | 0.61 | 0.56 | 0.06 |
| std | 0.09 | 0.15 | 0.13 | 0.04 | 0.03 |

**Table 3.**
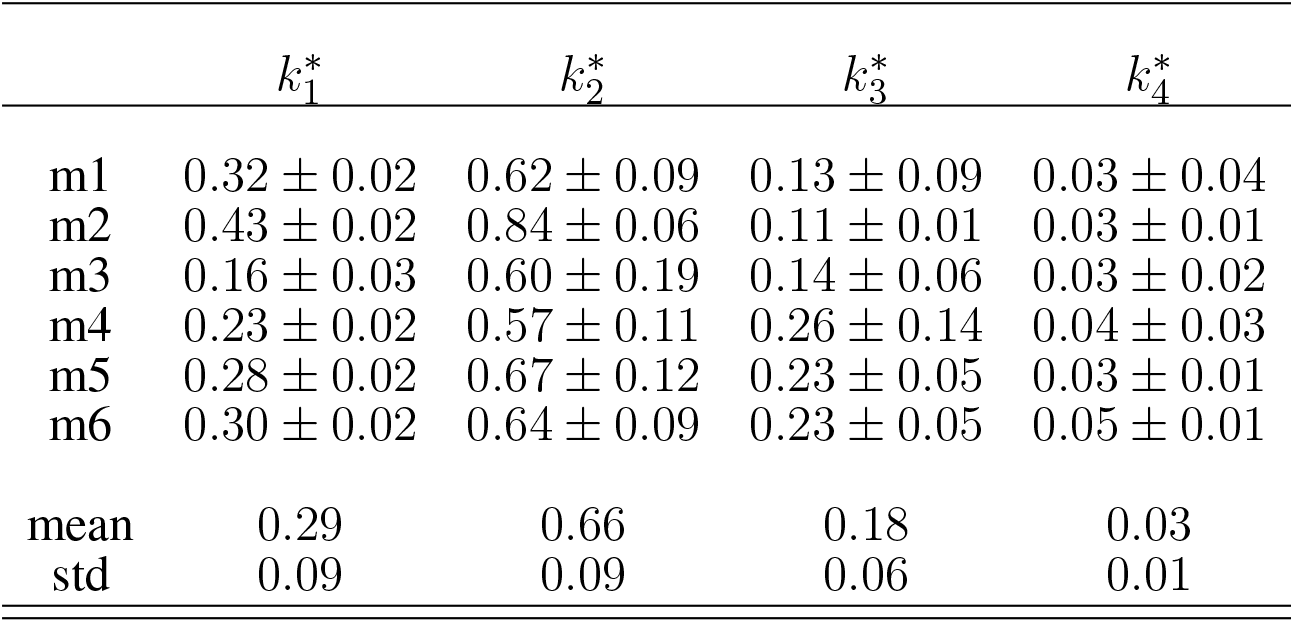
Reconstructed kinetic parameters (1/min) by the use of the SCM for the CT26 tumor tissue of the FDG–PET experimental group of six mice, as mean and standard deviation over 50 runs of the reg-GN algorithm. The last two lines report mean and standard deviation of each kinetic parameter computed over the mean estimates of the six murine experiments.

| | $k_1^*$ | $k_2^*$ | $k_3^*$ | $k_4^*$ |
| --- | --- | --- | --- | --- |
| m1 | $0.32 \pm 0.02$ | $0.62 \pm 0.09$ | $0.13 \pm 0.09$ | $0.03 \pm 0.04$ |
| m2 | $0.43 \pm 0.02$ | $0.84 \pm 0.06$ | $0.11 \pm 0.01$ | $0.03 \pm 0.01$ |
| m3 | $0.16 \pm 0.03$ | $0.60 \pm 0.19$ | $0.14 \pm 0.06$ | $0.03 \pm 0.02$ |
| m4 | $0.23 \pm 0.02$ | $0.57 \pm 0.11$ | $0.26 \pm 0.14$ | $0.04 \pm 0.03$ |
| m5 | $0.28 \pm 0.02$ | $0.67 \pm 0.12$ | $0.23 \pm 0.05$ | $0.03 \pm 0.01$ |
| m6 | $0.30 \pm 0.02$ | $0.64 \pm 0.09$ | $0.23 \pm 0.05$ | $0.05 \pm 0.01$ |
| mean | 0.29 | 0.66 | 0.18 | 0.03 |
| std | 0.09 | 0.09 | 0.06 | 0.01 |

Using the reconstructed average values of the kinetic parameters it is straightforward to compute the solution of the Cauchy problem associated to BCM and SCM and therefore the concentrations in all functional compartments. Figure 4 refers to mouse m1 and illustrates these concentrations in the case of the free compartment, the phosphorylated compartment and the reticulum compartment for BCM and in the case of the free and phosphorylated compartments for SCM.

**Figure 4.**
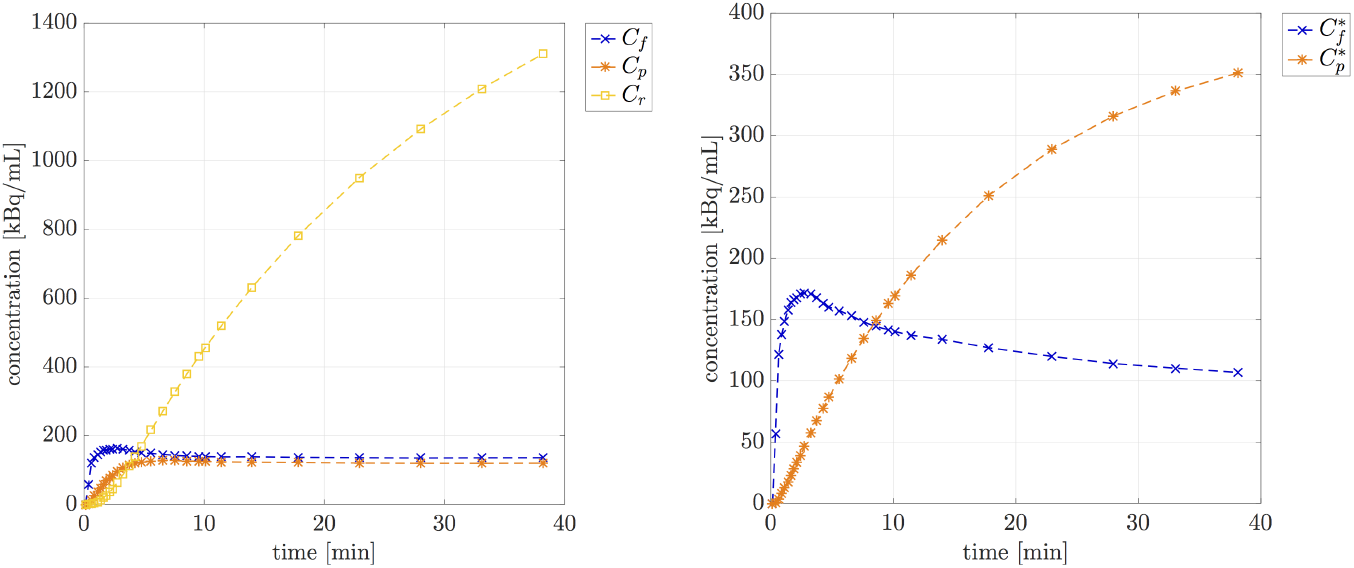
Model-predicted time curves of the compartment concentrations for mouse m1, computed by means of the reconstructed average values of the kinetic parameters: (a) *C*_*f*_, *C*_*p*_ and *C*_*r*_ of BCM; (b) 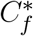, and 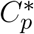 of SCM.

## 5 DISCUSSION OF RESULTS

The reconstructed values of the kinetic parameters in Table 2 and Table 3 are rather stable with respect to the murine models considered, which seems to imply that these numbers describe characteristic kinetic properties of FDG inside the CT26 tissue, independently of the specific murine experiment. Further the results of the analysis based on SCM are coherent with the ones obtained in the literature in the case of other kinds of carcinoma, for both mice and humans (Røe et al., 2010) (Rusten et al., 2013). This also confirms the reliability of the numerical method applied for model reduction.

Considering now mouse m1 as representative of all FDG-PET CT26-tissue experiments, Figure 4 shows that the concentrations of free tracer predicted by the two models are very close; further, in both cases FDG accumulation in its own accumulation pool (ER for BCM, cytosol for SCM) occurs with growing concentration (the concentration in ER is four times bigger in absolute values just because ER is a small fraction of the cytosolic phosphorylated compartment).

The numbers in Table 2 and Table 3 quantitatively compare the performances of the two compartmental models while explaining the same experimental data. In both cases the inversion procedure relies on both the model differential equations (equation (1) for BCM) and the measurement equations, i.e. the equations connecting the models to the data (equation (14) for BCM). The measurement equations are highly influenced by the volume fractions defining the amount of total volume occupied by each specific compartment. These volume fractions are different in BCM and SCM, i.e. the same compartment in the two models occupies a different relative volume. Given all this, the results presented in the two tables seem rather coherent. For example, *k*_1_ and 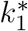 are statistically equivalent and the same occurs for *k*_6_ and 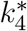: the reason for that is that these rate constants reflect the input process from the blood and the output process of dephosphorylation, respectively, and these two processes are just mildly dependent on the model adopted. On the other hand, the discrepancies between *k*_2_ and 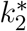, as well the ones between *k*_3_ and 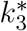 are due to the different volume fractions utilized in the two models, which reflects a different kinetics described by the two systems. Anyhow, it should be noticed that the estimated value of *k*_3_ obtained from direct measurements of the Michaelis-Menten parameters for the phosphorylation rate (Muzi et al., 2001) is about 0.9 (1/min), which is significantly closer to BCM kinetics rather than SCM kinetics. Therefore the two models provide a comparable data fitting but BCM seems more consistent with biochemistry measurements.

## 6 CONCLUSIONS

The main result of the present study is that, according to the reconstruction provided by BCM, FDG accumulates in phosphorylated form in ER in the same way this happens in cell cultures. This conclusion arises from an analysis where, in principle, the two available pools for phosphorylated tracer, cytosolic and ER-localized, are treated on the same level, in that no *a priori* constraints are imposed to the model. It should be noted that this result has been found in an experimental setting which is highly different with respect to the one considered in (Scussolini et al., 2019): (i) cell cultures and tissues have been inserted in different environments (clean incubation medium vs heterogeneous background, including blood and interstitial tissue), are constituted by different type of cancer cells (4T1 vs CT26), and occupy different total volumes; (ii) the datum of tracer consumption has been obtained through processes based on direct measurements of the emitted radiation (LT signals) and analysis of reconstructed images of radioactivity distribution (PET images); (iii) the IF of the cell system is almost constant, while the IF of the tissue system shows a sharp peak at the initial time. However, despite all these discrepancies, reconstructed FDG kinetics has shown rather stable characteristics with respect to the biological systems considered, and the similar performance of the BC model in such dissimilar environments is a further strong indication of its reliability, as mathematically confirmed by the identifiability considerations given in Section 3 of the present paper.

The results provided by the application of BCM need for further investigations. In fact, the basic scheme of this approach is rather flexible and may be modified to allow for consideration of peculiarities of specific organs, as done for classical models in (Garbarino et al., 2014, 2015), may be associated with reference tissue formulations (see (Scussolini et al., 2018) and references cited therein), or to pixel-wise analysis (see (Scussolini et al., 2017) and references cited therein).

